# A knowledge-based system for personalised lifestyle recommendations: Design and simulation of potential effectiveness on the UK Biobank data

**DOI:** 10.1101/2022.12.02.518736

**Authors:** Francesca Romana Cavallo, Christofer Toumazou

## Abstract

Mobile health applications, which employ wireless technology for healthcare, can aid behaviour change and subsequently improve health outcomes. Mobile health applications have been developed to increase physical activity, but are rarely grounded on behavioural theory and employ simple techniques for personalisation, which has been proven effective in promoting behaviour change. In this work, we propose a theoretically driven and personalised behavioural intervention delivered through an adaptive knowledge-based system. The behavioural system design is guided by the Behavioural Change Wheel and the Capability-Opportunity-Motivation behavioural model. The system exploits the ever-increasing availability of health data from wearable devices, point-of-care tests and consumer genetic tests to issue highly personalised physical activity and sedentary behaviour recommendations. To provide the personalised recommendations, the system firstly classifies the user into one of four diabetes clusters based on their cardiometabolic profile. Secondly, it recommends activity levels based on their genotype and past activity history, and finally, it presents the user with their current risk of developing cardiovascular disease. In addition, leptin, a hormone involved in metabolism, is included as a feedback biosignal to personalise the recommendations further. As a case study, we designed and demonstrated the system on people with type 2 diabetes, since it is a chronic condition often managed through lifestyle changes, such as physical activity increase and sedentary behaviour reduction. We trained and simulated the system using data from diabetic participants of the UK Biobank, a large-scale clinical database, and demonstrate that the system could help increase activity over time. These results warrant a real-life implementation of the system, which we aim to evaluate through human intervention.

## I. Introduction

mHealth (mobile health) technology is the use of wireless technology in medical care that can improve health outcomes due to its widespread appeal, accessibility and ability to reach large populations at a low-cost [1]. Many mHealth interventions have been developed to promote physical activity (PA), which mainly use wrist-worn or smartphone accelerometers and employ simple digital techniques. These mHealth interventions rarely utilise a holistic approach in attempting to positively alter one’s behaviour, such as grounding the intervention on behavioural theory and personalisation, which has been positively correlated with successful behaviour changes [2]. A systematic review and meta-regression [3] found that personalisation is effective in changing lifestyle behaviours and that the source of data used in the personalisation greatly affects the effectiveness of the intervention. Therefore, one of the main research gaps in behaviour change interventions is the effective use of personalisation, which can increase engagement and effectiveness of the interventions by improving one’s perceived skills and motivation [4]. According to the Self-Efficacy Theory [5], motivation is driven by self-efficacy, i.e. a person’s belief in their ability to succeed, which is developed through repeated successes to challenges that become more difficult as the skills improve. Consequently, personalisation of the intervention is critical to ensure that the subject builds high self-efficacy to succeed in the behaviour change process.

To achieve personalisation, interventions using artificial intelligence, specifically recommender or expert systems, have been developed. For example, Ni et al. [6] used a long short-term memory recurrent neural network (RNN) trained on user identity, sport type and historical workout sequences to recommend workout routes based on personal criteria (such as workout length and heart rate) and expectations. The model also predicts whether a user’s heart rate will exceed a user-set threshold if they continue at the current pace and suggest ways to hit the user’s target. Mahyari et al. [7] developed an intervention with RNN personalised such that the recommendations are highly likely to be performed based on past exercise history. Zhao et al. [8] developed and tested the efficacy of a tree-like personalised gamification intervention that recommends activity type, time and location based on the user’s general information, personality, and daily activity data from smartphone or wearable tracker. However, these works make limited use of personalisation compared to the levels that could be achieved given the ever-increasing availability of personal data. In recent years, and more specifically with the COVID-19 pandemic, the availability of point-of-care [9] and consumer genetic tests [10] offers the possibility of collecting even more health-related data to develop technology-driven interventions. For example, DnaNudge, a direct-to-consumer genetic service informed by behavioural science, developed an expert system that recommends food products with a suitable nutritional value depending on one’s genetic risk of developing metabolic conditions [11]. DnaNudge also encourages users to reduce sedentary time, with the penalty of having decreased food choices until a preset number of steps is achieved. However, while DnaNudge uses genetic information to recommend food choices, it does not use genotypes specific to physical activity, nor does it provide activity recommendations; rather, it lets the user set their preferred activity levels.

The knowledge-based system presented in the paper addresses the gaps in the current literature on recommender systems for PA. To demonstrate the concept, the system is developed and evaluated on a type 2 diabetes (T2D) population. T2D is a highly prevalent chronic condition linked to insulin resistance caused by obesity, a diet high in carbohydrates and inactivity, and is managed through lifestyle changes and drug therapy. Lifestyle management, including PA promotion, is the first-line therapy to improve glucose control in T2D [12]. In addition, with the recent understanding of the damages of sedentary behaviour (SB)—i.e., waking activities that require lying or sitting—people with T2D are now also recommended to reduce their sedentary time, as this is associated with increased glucose metabolism biomarkers [13]. To promote a healthier lifestyle effectively, behaviour-change strategies are recommended. In particular, technology-enabled interventions can be especially useful in increasing activity and reducing sedentary time, thus improving glucose control in T2D patients [14].

The present system aims to enable the behaviour change process by using several layers of personalisation: firstly by classifying the user into one of four diabetes clusters, secondly by recommending activity levels based on their genotype and past activity history, and finally by presenting the user with their risk of developing cardiovascular disease (CVD) based on their current activity levels. Another layer of personalisation is achieved by including leptin - monitored through a novel point-of-care system [15]—as a feedback biosignal to inform on the efficacy of the intervention towards weight loss. In fact, leptin is a biomarker for obesity management therapies as leptin resistance is related to obesity and leptin levels vary with dieting regimens [16]. To our knowledge, no previous system or intervention has used blood chemistry as a feedback signal for behavioural change interventions. Biomarkers have previously been used to assess the efficacy of interventions, but never as an integral part of the recommendation systems. Finally, the proposed system includes SB alongside PA. Previous works have overlooked SB, despite it being paramount to improving glucose control in T2D populations.

This paper presents the design and simulation of a knowledge-based system for personalised activity recommendations. First, we outline the system’s design as a behavioural intervention, implemented with the Capability-Opportunity-Motivation Behavioural model and the Behaviour Change Wheel. Secondly, we discuss the clustering of diabetes patients based on cardiometabolic biomarkers and their associations with PA and SB. Thirdly, we present the main components forming the proposed system, whose high-level diagram is shown in Fig. 1:

1. A classifier to categorise new patients in clusters of T2D. The optimal classifier was chosen to be a Support Vector Classifier, which achieved better accuracy than K-Nearest Neighbours and Random Forests;
2. The optimised recommendation algorithm, which uses past activity levels, leptin levels, and a polygenic risk score to issue the personalised recommendation;
3. A survival model that calculates the risk of developing CVD based on the patient’s activity levels. Several models for survival prediction were tested, and the Cox Proportional Hazards model was chosen due to its higher performance.

**Fig. 1:**
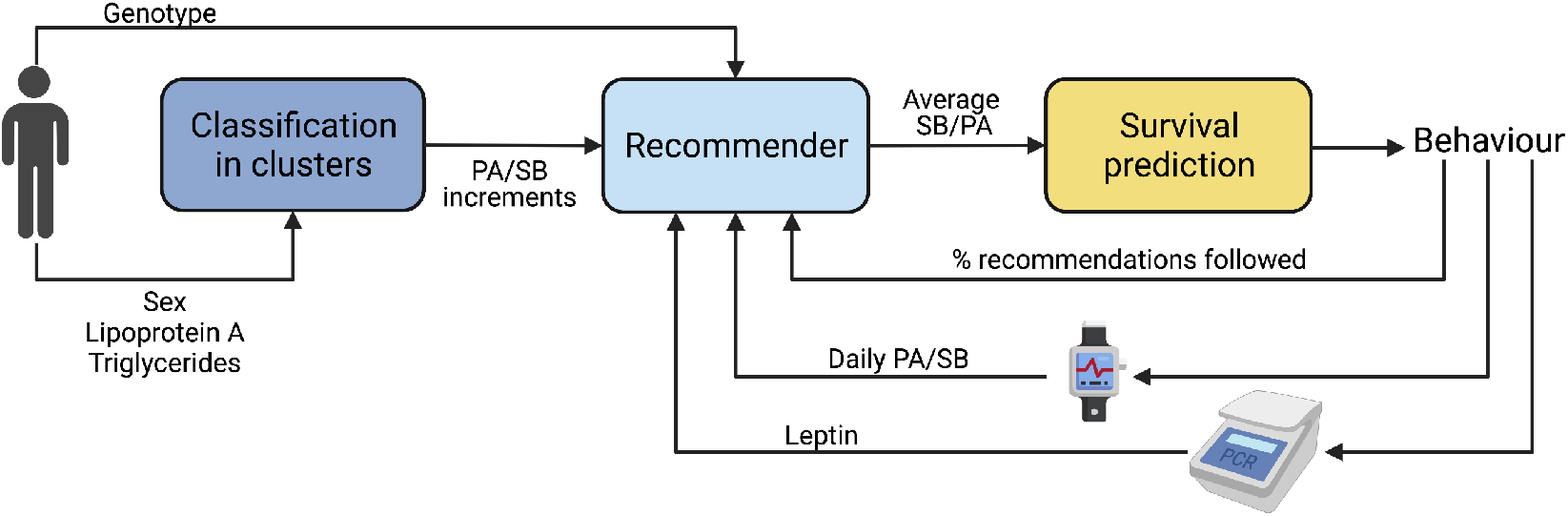
High-level diagram of the proposed system.

The recommendation system is designed as a content-based system[17], since it employs the user’s feedback to recommend activity levels. Moreover, the system employs both categorisation (by using cluster-specific parameters) and personalisation (by updating the features based on user-specific data). Finally, we present the results of simulations of the proposed system on the data from the UK Biobank, a large-scale clinical database with biomedical data on more than 500,000 participants.

Despite being trained and evaluated on a T2D population, the present system can be easily adapted to different populations with a desire to achieve a more healthy lifestyle by increasing physical activity and decreasing sedentary behaviour.

## II. Design of the system as a theoretically driven behavioural intervention

Despite the increasing complexity and number of mHealth interventions available, these technologies fail to induce long-term changes [1]. Reviews have highlighted the need to use behavioural theories to design interventions in order to achieve higher efficacy [2]. Therefore, we designed the high-level structure of the system following the Capability-Opportunity-Motivation Behavioural (COM-B) model and the Behaviour Change Wheel (BCW) [18]. The BCW provides a framework to develop complex interventions that is comprehensive, coherent and linked to a behavioural model, the COM-B model. This framework has been used to develop digital interventions that are both accepted by users and effective for a variety of behaviours, including improving diet [19], [20], [21], increasing physical activity [19], [22], [23] and reducing sedentary behaviour [24], management of hypertension [25] and diabetes [26], [27], smoking cessation [28], [29], alcohol reduction [30] and medication adherence [31], [32] or discontinuation [33]. The steps needed to develop an intervention following the BCW are: 1) Specifying the target behaviour; 2) identifying theoretical domains that explain the behaviour; 3) identifying how to target the theoretical domains; 4) selecting behaviour change techniques.

To design the system, we followed the steps mentioned above. Firstly, we specified the target behaviour (i.e. increasing PA and reducing SB) by identifying the factors associated through a literature review. Ovid and Embase databases were systematically searched (see the Supplementary Information for the search term) to include studies identifying barriers and facilitators to the target behaviours, specifically in type 2 diabetes patients. Five studies were selected [34], [35], [36], [37], [38] (a summary of the selected papers is provided in Supplementary Table S1) that identified factors associated with the target behaviours pertaining to all COM-B model domains (capability, opportunity and motivation). All factors associated with the target behaviours that were identified by the selected papers are presented in Table I, including those that cannot be addressed by a mHealth intervention (such as environmental factors), in order to have a more complete picture of the elements influencing physical behaviours in people with type 2 diabetes.

**TABLE I:**
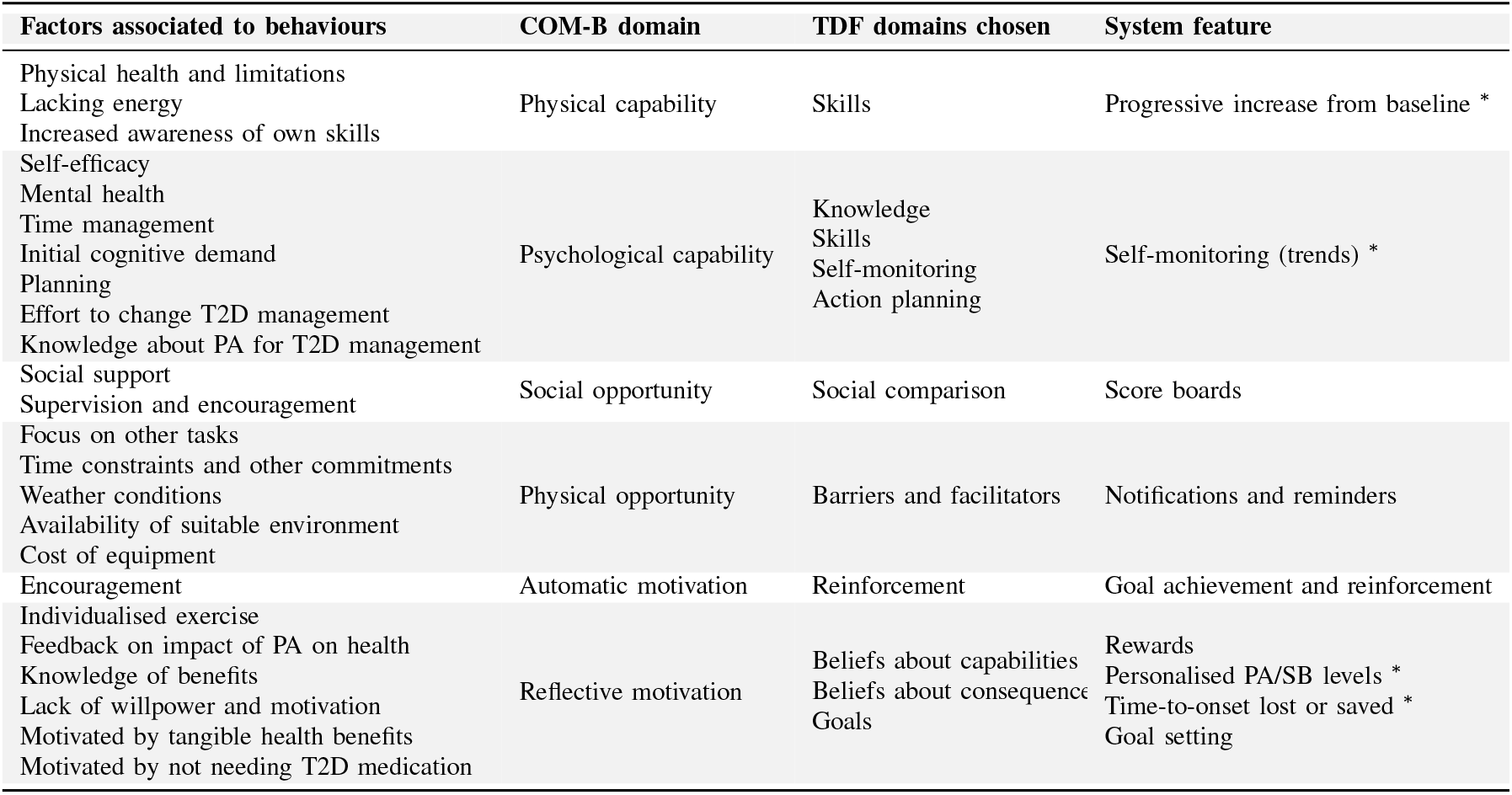
Implementations of the BCW for the design of a knowledge-based system for personalised PA and SB recommendations. Entries marked with (*) have been implemented in this work.

Secondly, we completed step 2 of the BWC by mapping the identified factors to the COM-B model. Finally, to complete steps 3 and 4, we identified behaviour change techniques through the help of the Theoretical Domains Framework (TDF) [39], and we mapped them to system features. The design process is shown in Table I.

According to the COM-B model, an intervention must match the patient’s abilities to be effective and sustained. Challenges recommended by the system have to align with the patients’ skills and constantly adjust as these change over time. Therefore, in the proposed system, the recommendations are initialised to the patient’s baseline activity and optimised over time depending on the probability of compliance and leptin levels. The probability of compliance with the recommendations is paramount to prevent feelings of failure in the patient, which can hinder engagement with the intervention. Moreover, PA and SB trends enable self-monitoring and self-regulation, which increase motivation. Finally, rewards are embedded in the system as a time-to-onset of comorbidities gained or lost in response to their new activity behaviour.

## III. Clusters of type 2 diabetes and associations with physical activity and sedentary behaviour

Given the high heterogeneity of T2D, several studies have explored the clustering of diabetic patients [40], [41], [42], [43], [44], [45], [46], and all have analysed the clusters in terms of disease risk and progression. The clusters were determined with various methods (K-means[41], [43], hierarchical clustering[42], [44], [45], TwoStep clustering[40], [46]) and using features including blood chemistry, anthropometric data and measures of glucose metabolism. However, none of the studies in the literature considered the associations between clusters and lifestyle risk factors. Determining such associations is essential to develop personalised behavioural therapies for T2D patients to lower the risk of comorbidities related to a poor lifestyle. Therefore, the first aim of this work is to quantify the differences in PA and SB between clusters and determine their effects on the risk of developing CVD, a condition commonly associated with T2D.

The sample used in the analyses was selected among the participants to the UK Biobank. The UK Biobank study design is described in detail in [47]. Briefly, the UK Biobank is a database of over 500,000 adults aged 37-73 recruited in the UK between 2006 and 2010. Ethics approval for the UK Biobank study was obtained from the North West Centre for Research Ethics Committee (REC reference: 21/NW/0157). Informed consent for the UK Biobank study was obtained from participants during the baseline assessment. The work described here was approved by the UK Biobank under application ID 56132. All analyses were performed in accordance with the relevant guidelines and regulations.

The workflow of the analyses conducted on the UK Biobank is described in Figure 2.

**Fig. 2:**
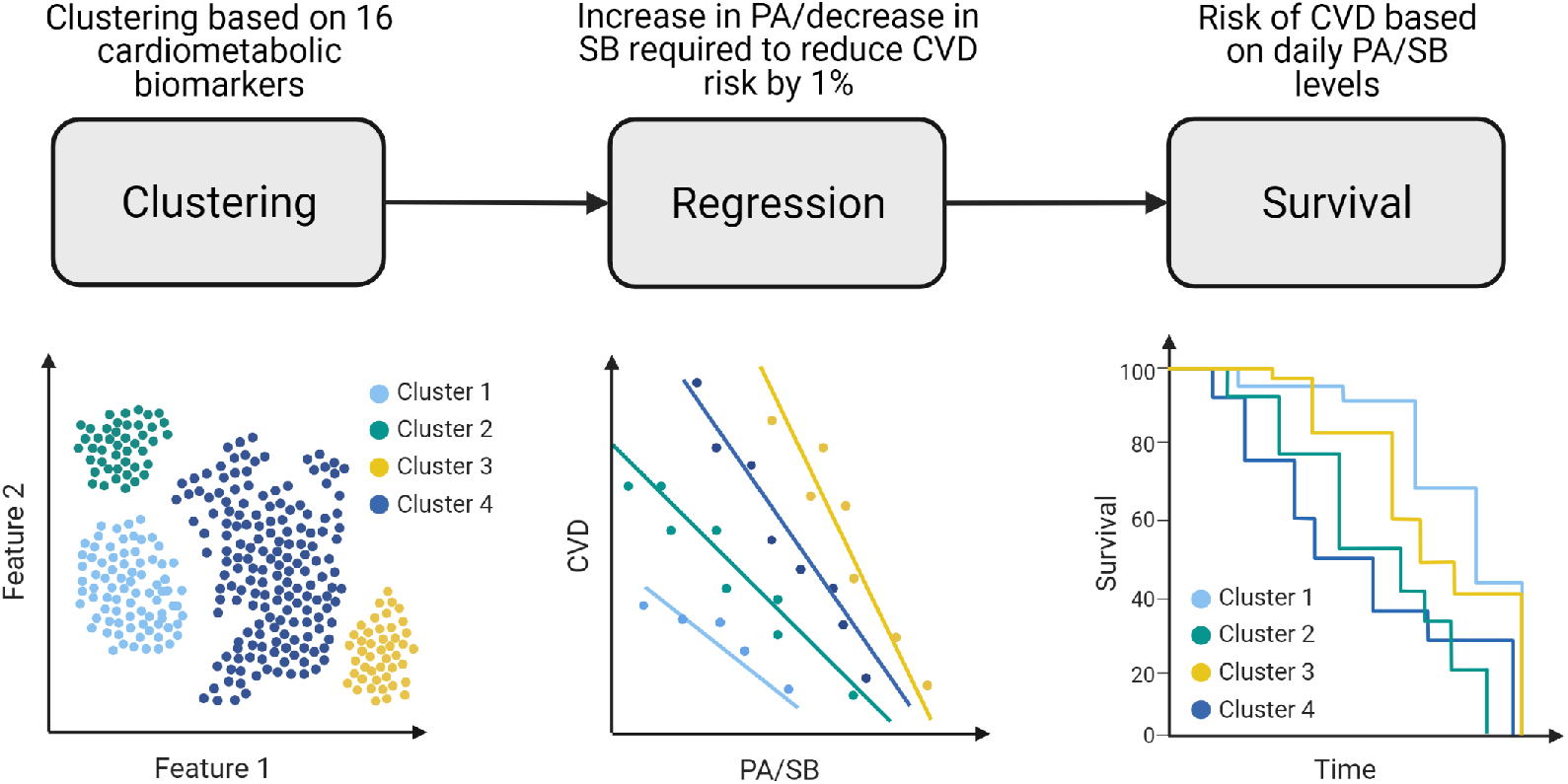
Flow diagrams illustrating the simulations performed on the UK Biobank dataset. The graphs do not depict real data, but are for visualisation only.

### A. Cluster analysis

Participants of the UK Biobank with a self-reported T2D diagnosis were selected. We selected 16 cardiometabolic markers measured at baseline: systolic blood pressure, diastolic blood pressure, pulse rate, age at diagnosis, waist circumference (WC), body mass index (BMI), apolipoprotein A, apolipoprotein B, C-reactive protein, cholesterol, HDL-cholesterol, LDL direct, glucose, HbA1c, lipoprotein A and triglycerides. Clustering was done using the markers rescaled to the [0,1] range. The dataset was split based on gender and analysed separately to avoid sex-dependent differences [40]. After removing participants with missing values for one or more biomarkers, 14,149 participants (9,054 males and 5,095 females) were included in the cluster analysis.

Four clustering methods were tested with the scikit-learn [48] and scikit-learn-extra in Python: k-means, k-medoids, agglomerative hierarchical clustering and Gaussian mixture models (GMM). The hyperparameters were tuned based on the silhouette width [49] metric. Cluster stability was evaluated using the bootstrapped Jaccard coefficient in the R fpc package [50], which computes the similarity between clusters on the whole sample to clusters on the bootstrapped dataset (1000 resamples). A cluster with a Jaccard coefficient value ≥ 0.75 is considered stable. The clustering methods were evaluated based on the silhouette, Calisnki-Harabasz and Davies-Boulding scores, and GMM with *n* = 4 and a spherical covariance matrix was chosen. The four clusters generated by GMM achieved a Jaccard score ≥ 0.75 and therefore were deemed stable. Differences in cardiometabolic biomarkers between clusters were assessed with one-way ANOVA (effect sizes reported as Cohen’s *d*), and a Bonferroni correction was applied to the *p*-value to account for multiple tests.

The cluster characteristics are summarised in Table II. Cluster 1 was characterised by lower T2D onset age, lower BMI/WC and improved cardiometabolic profile. Cluster 2 had an average T2D onset age (around 51 years for both males and females), higher BMI and slightly worse blood pressure and CRP than cluster 1. Cluster 3 and 4 were characterised by higher onset age and worse cardiometabolic profile, with cluster 4 having the highest values of blood pressure, lipids (combining cholesterol, HDL-C, LDL-C, apolipoprotein A and B, and lipoprotein A), glucose, HbA1c and CRP across all clusters. The differences between clusters were similar in males and females, as shown in Table II. However, the prevalence of comorbidities differed between clusters, with males’ cluster 3 having the highest prevalence of hypertension and CVD (31% and 65%, respectively) and males’ cluster 4 having the highest prevalence of high cholesterol (40%). For females, cluster 4 had the highest prevalence of high cholesterol (43%) and CVD (25%); and cluster 3 had the highest prevalence of hypertension (67%). Cluster 1, for both males and females, was characterised by significantly higher insulin use (28% in males, 33% in females vs < 19% in other clusters).

**TABLE II:** Summary of cluster characteristics. Biomarkers are marked as ‘better’ or ‘worse’ if the Cohen’s *d* in t-tests between the cluster and the whole sample (stratified by sex) was ≥ 0.25. (CVD: cardiovascular disease, HT: hypertension, HC: high cholesterol)

| *Males* | **Cluster 1** | **Cluster 2** | **Cluster 3** | **Cluster 4** |
| --- | --- | --- | --- | --- |
| **Onset Age** | Average | Average | Average | Average |
| **BMI/WC** | Better | Average* | Average* | Worse* |
| **BP** | Average* | Average* | Average* | Average* |
| **Cholesterol** | Better | Better | Worse* | Worse* |
| **Triglycerides** | Better | Better | Worse* | Worse* |
| **Glucose** | Average* | Average* | Average* | Average* |
| **HbA1c** | Average* | Average* | Average* | Average* |
| **CRP** | Average* | Average* | Average* | Average* |
| **Highest odds** | Insulin use |  | HT, CVD | HC |
| *Females* |  |  |  |  |
| **Onset Age** | Lower | Average | Average | Average |
| **BMI/WC** | Better* | Average* | Average* | Worse* |
| **BP** | Average | Average* | Average* | Average* |
| **Cholesterol** | Better | Average* | Average* | Worse* |
| **Triglycerides** | Better | Better | Worse* | Worse* |
| **Glucose** | Average* | Average* | Average* | Average* |
| **HbA1c** | Average* | Average* | Average* | Average* |
| **CRP** | Average* | Average* | Average* | Average* |
| **Highest odds** | Insulin use |  | HT | HC, CVD |

\*Biomarker values beyond the normal range, as indicated in the supplementary information.

### B. Regression analysis

Logistic regression was done with statsmodels [51] to assess inter-cluster differences in associations between PA (or SB) and CVD. A full list of illnesses classified as CVD, including hypertension and high cholesterol, is included in the supplementary information. After removing participants with missing accelerometer data, or without at least 3 days of data and with data in each one-hour period of the 24-hour cycle (scattered over multiple days), *n* = 1805 participants were included in the regression analysis. Firstly, we classified the accelerometer data into SB, light PA, walking, moderate-to-vigorous PA (MVPA) and sleep with balanced random forests and Hidden Markov models (with the accelerometer package by [52]). Secondly, we partitioned the day into time spent in overall PA (including time spent in light PA, walking and MVPA), SB and sleep, and we included the accelerometer in the regression model with a compositional approach [53]. In a compositional framework, time spent in the three different activities is considered as a relative proportion of the overall time budget (24 hours), such that the vector 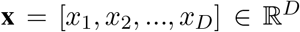, with *D* = number of activities and with 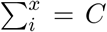, is constrained by the closure constant *C* = 24 hours. The closure constant implies multicollinearity among the activities and thus conventional statistical methods cannot be employed with compositional data [54]. The isometric log-ratio (ILR) transformation maps the data from the constrained simplex space to the unconstrained real space, which allows for the application of regression. Therefore, the transformation *z* = *ILR*(*x*) is applied to the accelerometer data as follows:

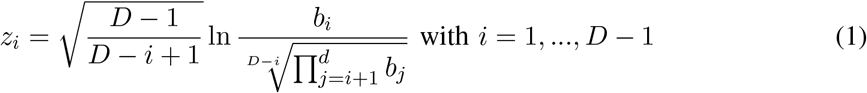

In this paper we report only regression coefficients for *z*_1_, since the regression coefficient *β*_1_ for the first *ILR* coordinate *z*_1_ represents the strength of the association between the chosen activity and the outcome, while *z*_1+*i*_ cannot be interpreted in a meaningful way. The *ILR* regression coefficients can be used to estimate the minutes of activity required to reduce the log(odds) of CVD, as Dumuid et al. [54] proved that for a 3-activity composition the following holds:

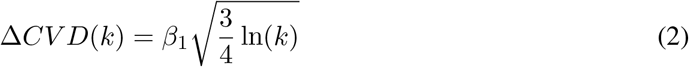

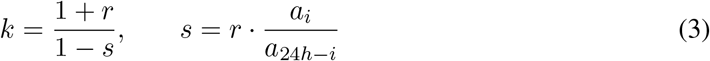

with *k* being change in daily activity composition, *r* being the change in primary activity *a_i_* compared to the sample mean and *s* being the corresponding change in the remaining activity components *a*_24*h*–*i*_. The odd of CVD and mean time spent in PA and SB for each cluster is presented in Table III. The time steps required to reduce the risk of CVD by 1% for each cluster are presented in Table IV. Some of the steps in Table IV were adjusted, given the wrong step direction due to *p*-value> 0.05. Specifically, the PA time step for clusters 2 ad 4 are replaced by the time step for the whole sample (20 minutes/day). The SB time step for cluster 2 are set to 18 minutes/day, given the similarity in SB levels with cluster 1 and 3.

**TABLE III:** Odds of CVD and geometric mean of time spent in physical activity by cluster.

| Cluster | Ill ($n$) | Healthy ($n$) | CVD odds | PA mins | SB mins |
| --- | --- | --- | --- | --- | --- |
| Whole sample | 1264 | 351 | 3.6 | 243 | 509 |
| 1 | 391 | 138 | 2.8 | 262 | 487 |
| 2 | 326 | 84 | 3.9 | 241 | 514 |
| 3 | 224 | 61 | 3.7 | 233 | 522 |
| 4 | 323 | 68 | 4.8 | 229 | 528 |

**TABLE IV:** Associations between CVD and compositional measures of physical activity and conversion from compositional coefficients to time (minutes/day) of physical activity and sedentary behaviour required to reduce the risk of CVD by 1%. The log(odds) correspond to changes in CVD log-odds for increases in time spent in the given behaviour and corresponding decrease in the others.

| PA | $n$ | log(odds) ($p$-value) | Step (mins/day) |
| --- | --- | --- | --- |
| Sample | 1805 | -0.41 (0.009) | 20 |
| Cluster 1 | 602 | -0.83 (0.001) | 8 |
| Cluster 2 | 445 | 0.06 (0.84) | -108 |
| Cluster 3 | 432 | -1.45 (0.0003) | 6 |
| Cluster 4 | 326 | 0.2 (0.55) | -47 |
| SB | $n$ | log(odds) ($p$-value) | Step (mins) |
| Sample | 1805 | 0.17 (0.44) | -72 |
| Cluster 1 | 602 | 0.51 (0.15) | -19 |
| Cluster 2 | 445 | -0.84 (0.07) | 16 |
| Cluster 3 | 432 | 0.74 (0.13) | -17 |
| Cluster 4 | 326 | 0.11 (0.82) | -141 |

The effect of common confounders including age, WC, alcohol intake, income and smoking status as covariates was examined and the final model was selected with backwards selection—only covariates reaching significance (*p*-value < 0.05) were retained in the final models. Since only WC, smoking status and age were significant covariates, participants with complete data for these variables were included in the analysis (*n* = 1805). Sensitivity analysis was done on a subset of participants with no missing data for all covariates (including alcohol intake and income) with *n* = 1615. Regression on the subset produced similar results to the original sample, thus confirming the robustness of the coefficients. Given the similar cluster characteristics (see Table II), males and females were combined. After backward selection, the final models for the whole sample are:

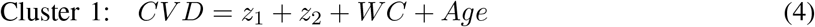

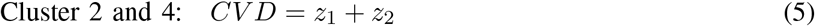

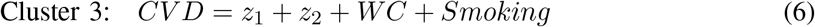

## IV. The recommendation system

### A. Classification into diabetes cluster

The first component of the proposed system is the classification of patients into the diabetes clusters. To select the best classifier, three models—K-nearest neighbours (KNN), support vector classifier (SVC) and random forest (RF)—were trained and validated on the clustering set (*n* = 26400). The test set was selected as the set of under-68 years old subjects (*n* = 348) later used in the prediction of survival times, as this set is used to simulate the system and thus should not be used to train the classifier.

For each of the three models, features were selected through forward sequential feature selection in mlxtend [55], using balanced accuracy as the selecting metric. Based on the best accuracy scores, the data were standardised and three features were selected for KNN and SVC (sex, lipoprotein A and triglycerides), while four were selected for RF (sex, pulse rate, LDL-cholesterol and triglycerides). The model hyperparameters were tuned with randomised search cross-validation, using the scikit-learn *RandomizedSearchCV* function which implements a fit and score method [48], with balanced accuracy being selected as the scoring method. The tuned models with the optimal number of features were then validated on the test set (see Table V). While the three classifiers achieved comparable performance, the support vector classifier was chosen as the preferred classifier as it uses fewer features than the Random Forest and reached a better accuracy than the KNN.

**TABLE V:** Classifier performance and number of features selected through sequential selection. (KNN: K-nearest neighbours, SVC: Support vector classification, RF: Random forest

| Classifier | N. of features | Training accuracy | Test accuracy |
| --- | --- | --- | --- |
| KNN | 3 | 0.98 | 0.99 |
| SVC | 3 | 0.99 | 0.994 |
| RF | 4 | 1 | 1 |

### B. Optimised recommendation module

The main component of the system is the recommendation module, which issues optimised recommendations to improve metabolic health while keeping the probability of compliance high. The core inputs for the recommendation module are leptin levels, probability of compliance, and upper and lower limit of PA and SB.

Leptin varies in relation to dieting therapies and thus, leptin levels are a real-time physiological signal providing feedback on the efficacy of the intervention. During diets, leptin levels change at a greater rate than weight and can predict weight loss two weeks in advance [56]. Therefore, monitoring leptin can be more meaningful than monitoring weight, and can be employed as feedback to adjust recommendations if future weight gain is predicted. Monitoring leptin levels is also relevant for chronic disease management, as hyperleptinemia is correlated to increased risk of obesity and cardiovascular outcomes in T2D patients [57].

The probability of compliance (Eq. 7) is related to the historical levels of activity and therefore, encodes the skill level of the patients. In order to keep compliance high, the recommendations are adjusted based on the proportion of recommendations followed in the past and their past activity levels. To include the effect of genetics on motivation, the recommendation score is also weighted by a polygenic risk score. The polygenic risk score is calculated using effect sizes reported by Doherty et al. [58] that quantify the association between selected Single Nucleotide Polymorphisms (SNP) and PA or SB. Information on the selected SNP is given in Table VI.

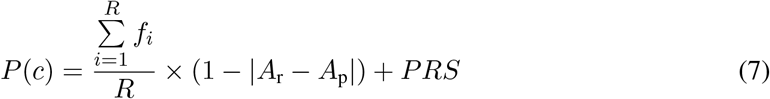

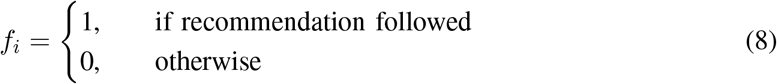

*R* is the number of recommendations issued and 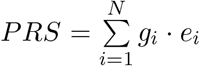 is the polygenic risk score, where *g_i_* is the genotype and *e_i_* is the effect size corresponding to SNP *i*. The factor *α* = 1 – | *A*_r_ – *A*_p_| represents the influence of past activity on the probability of compliance, where *A*_r_ is the new recommended activity and *A*_p_ is the predicted activity estimated through fitting a logistic function on the historical activity levels. To ensure 0 ≤ *α* ≤ 1, the recommended and predicted activity are calculated as daily proportions. The algorithm to determine the optimised recommendation is shown in Fig. 3.

**TABLE VI:** Genetic markers used in the polygenic risk score calculation. SNP: single nucleotide polymorphism, AF: allele frequency.

| Trait | SNP | Chromosome | Nearest Gene | Effect Allele | Alternative allele | AF |
| --- | --- | --- | --- | --- | --- | --- |
| PA | rs6775319 | 3 | SATB1-AS1 | A | T | 0.271 |
| PA | rs6895232 | 5 | LINC01470 | T | A | 0.663 |
| PA | rs564819152 | 10 | SKIDA1 | A | G | 0.679 |
| PA | rs2696625 | 17 | KANSL1-AS1 | A | G | 0.77 |
| PA | rs59499656 | 18 | SYT4 | A | T | 0.655 |
| SB | rs61776614 | 1 | FAAP20 | C | T | 0.925 |
| SB | rs1858242 | 3 | LOC105377146 | A | G | 0.259 |
| SB | rs26579 | 5 | MEF2C-AS2 | G | C | 0.415 |
| SB | rs25981 | 5 | EFNA5 | G | C | 0.531 |
| SB | rs6870096 | 5 | NMUR2 | C | G | 0.321 |
| SB | rs34858520 | 7 | CALN1 | A | G | 0.558 |

**Fig. 3:**
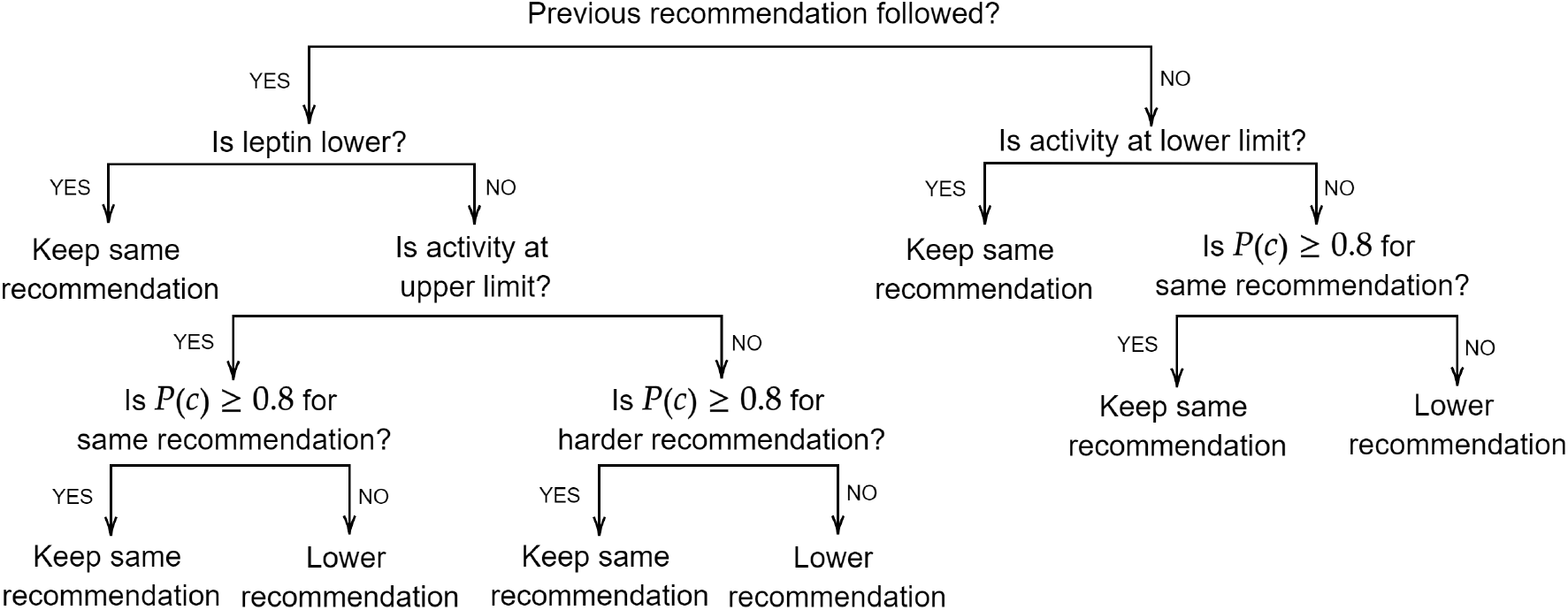
Diagram of the algorithm to generate the optimised recommendations using historical activity levels and the probability of compliance to the updated recommendation.

### C. Survival analysis for risk communication

To communicate the risk of developing CVD in response to one’s levels of PA and SB, the survival function of each cluster was estimated. In order to predict survival time from PA and SB levels simultaneously, activity data was not transformed with ILR transformation. The time between birth and onset of CVD was used as the time-to-event of interest, with the event being the diagnosis of CVD. The observations were censored at 68 years, and subjects that had not reached this age at the baseline visit were removed, leaving a sample size of *n* = 1215. Smoking, sex, age, triglycerides, diastolic blood pressure (DBP), total cholesterol, LDL-cholesterol and WC were included as covariates in the models, as they are the most common predictors for CVD [59]. After examination of the partial residual plots, a non-linear term 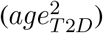 and two interaction terms (*PA × age* and *PA × DBP*) were included. All features were scaled to the [0, 1] range. Four models were compared with k-fold cross-validation (*k*=10, 100 iterations): Cox’s proportional hazards, Weibull AFT, Log-logistic and Log-normal AFT models. The analysis was done with the Python lifelines package [60]. The Cox’s model was selected as it obtained the highest concordance score (0.74 for Cox, Weibull and Log logit, 0.73 for Log normal), which measures the accuracy of the ranking of predicted times.

## V. Results

### A. Physical activity and leptin simulations

Physical activity was simulated based on whether the proposed recommendation was followed or not, which was modelled through different probability distributions (see next section). If the recommendation was followed, the PA (or SB) level is set equal to the recommendation, otherwise, it is set equal to the level recorded (or simulated) in the previous day.

Baseline leptin levels were predicted from gender, age, BMI, WC, arm fat mass, energy intake, PA and smoking using the regression coefficients estimated by Zuo et al. [61]. Missing data were replaced by the sample mean. However, for energy intake there was a significant amount of missing data, and values were replaced through multivariate sample imputation [48]. Leptin was updated based on the results found in a systematic review by Fedewa et al. [62], which found that engaging in chronic exercise (≥ 2 weeks) is associated with a reduction of leptin with an effect size of 0.24. Consequently, leptin is reduced by the effect size if the recommendations are followed for 2 weeks; otherwise, it is kept unchanged.

### B. Effect on behaviour change

The effectiveness of the system in inducing behaviour change was simulated on the *young* dataset with no missing genetic data (*n* = 290). The probability of following the recommendations on any given day was sampled from a Bernoulli distribution, and we modelled the parameter *p* with 3 different functions that reflect the time-dependent nature of adherence to human interventions. The models were estimated from data reported by Finkelstein et al. [63] (model in Eq. 9, Fig. 4A), who measured the percentage of participants who wore the activity tracker for at least 1 day in a week and by Meyer et al. [64] (model in Eq. 10, Fig. 4B), who recorded the average number of days in a week that participants wore the tracker. The parameters for both models were estimated with least squares [65]. The third model was time-invariant, with constant *p* over the duration of the intervention (Eq. 11).

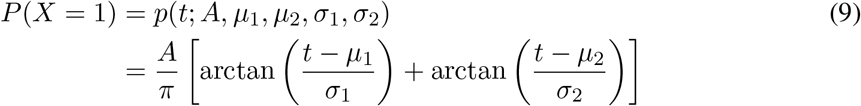

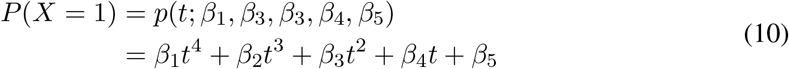

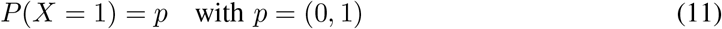

**Fig. 4:**
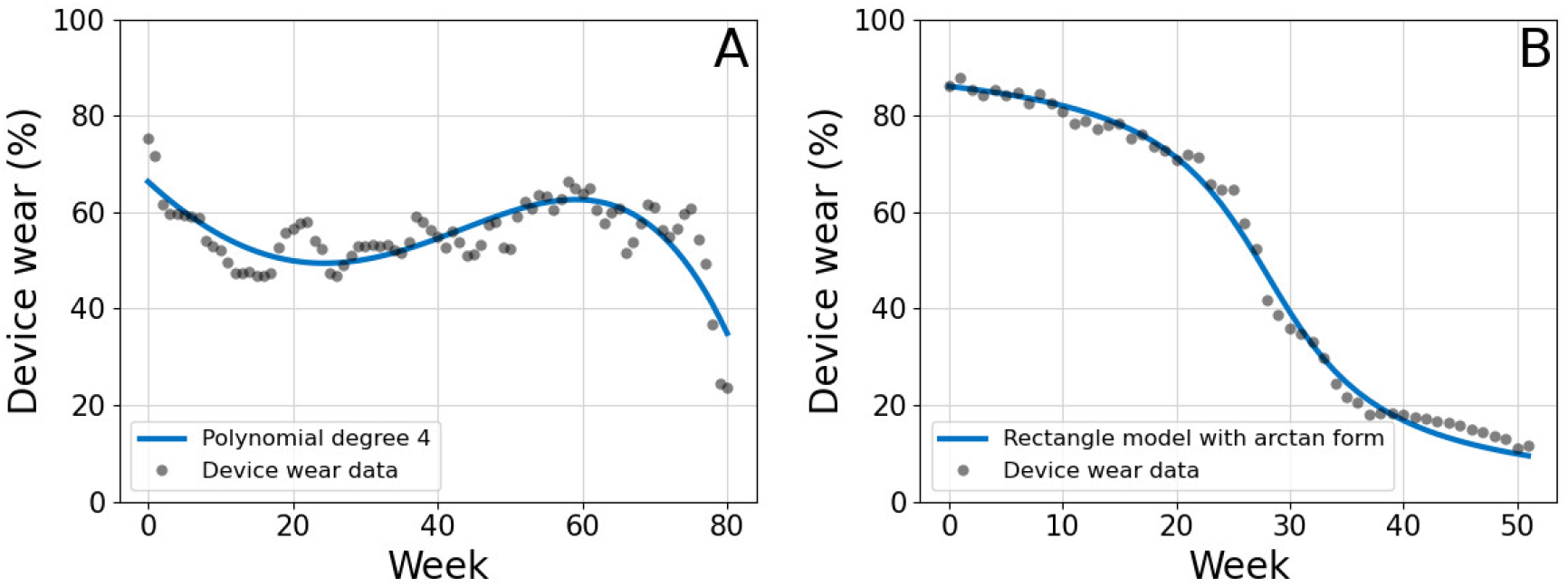
Progression of accelerometer device wear during time as reported by Finkelstein et al. [63] (A) and Meyer et al. [64] (B). The data was used to fit the models for adherence probability described in Eq. 9 and Eq. 10, respectively.

The proposed system was simulated with the aforementioned models for a duration of 365 days. The resulting activity and adherence profiles for the *young* dataset are presented in Fig. 5. Within the *young* dataset, four participants had follow-up accelerometer data available, collected on average approximately 1.5 years after the baseline measurements. Consequently, it was possible to compare the activity at follow-up with the simulated activity at a time matching the time difference between baseline and follow-up measurements. Fig. 6 shows that, although only a constantly high adherence (*p* = 0.8) to the intervention would bring significant improvements in activity compared to baseline, all adherence models achieved improvements compared to activity measured at followup. For PA, such difference was statistically significant for all models evaluated (*p*-value < 0.05, tested with paired t-test).

**Fig. 5:**
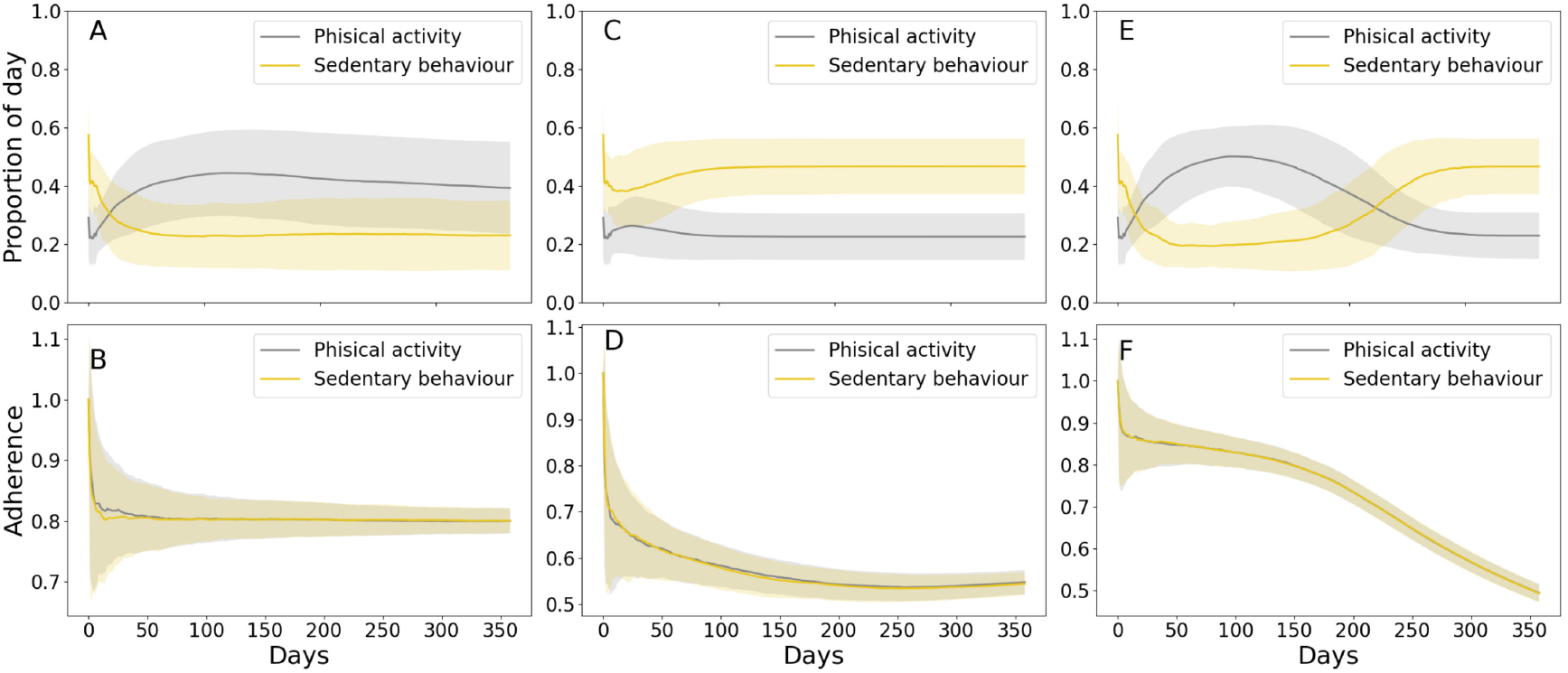
Simulated activity and adherence (i.e. proportion of recommendations followed) with timevarying probability according to Eq. 11 with *p* = 0.8 (A), Eq. 10 (B) and Eq. 9 (C). The top graphs show the proportion of the day spent in physical activity and sedentary behaviour across the simulated intervention. The bottom graphs show the simulated adherence to the intervention as a function of days. Data presented as mean and standard deviation.

**Fig. 6:**
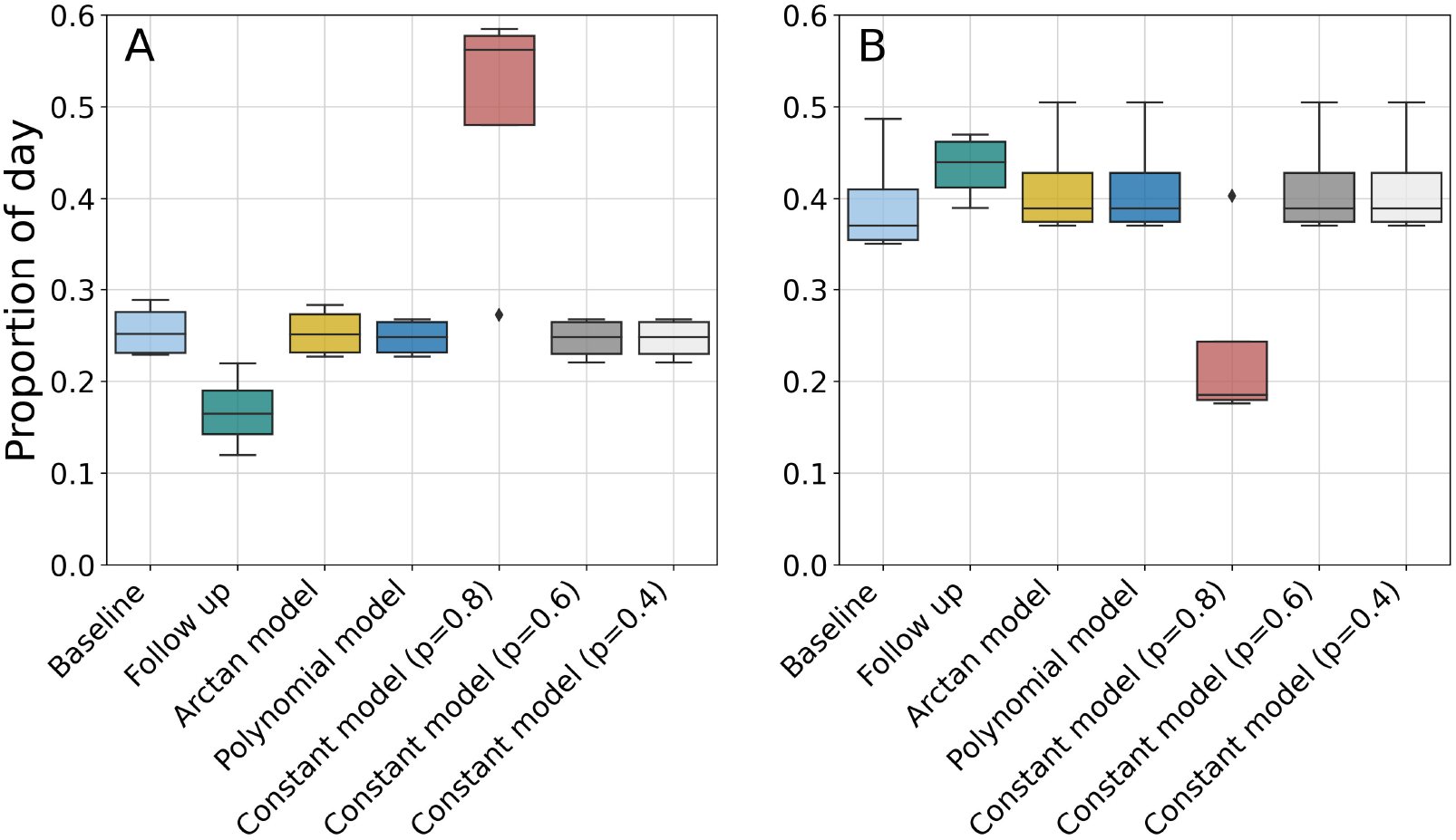
Comparison between simulated physical activity (A) and sedentary behaviour (B) after one year of personalised intervention and real-life measured activity at baseline and follow-up.

### C. Effect of physical activity and sedentary behaviour on survival times

Fig. 7 shows the Kaplan Meier survival baseline functions estimated on censored subjects (*n* = 1215), where the biggest difference is observed between clusters 1 and 4. This result is expected, since cluster 4 presents a worse cardiometabolic profile than cluster 1 and thus, is more prone to CVD development. Fig. 8 shows the effects of changing SB and PA on cluster 2 as a representative example; the effect of PA is visibly pronounced, while the effects of varying sedentary time are more subtle. Median survival times for the clusters, calculated as the month at which the survival function crosses *p* = 0.5, are shown in Table VII. They were predicted for *young* subjects, i.e. younger than 68 years (*n* = 348) that were not included in the baseline model estimation.

**Fig. 7:**
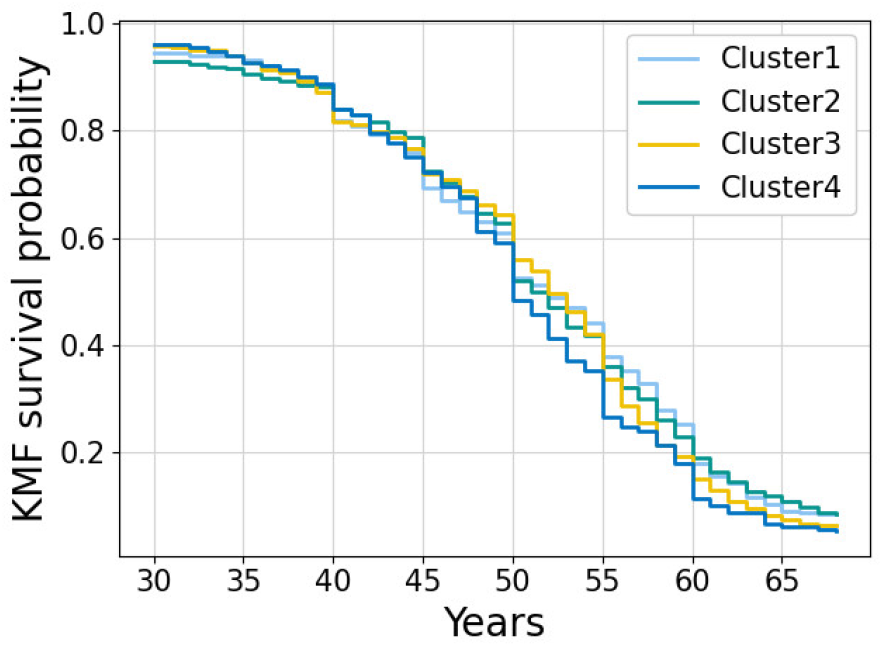
Kaplan-Meier survival probability for the four clusters.

**Fig. 8:**
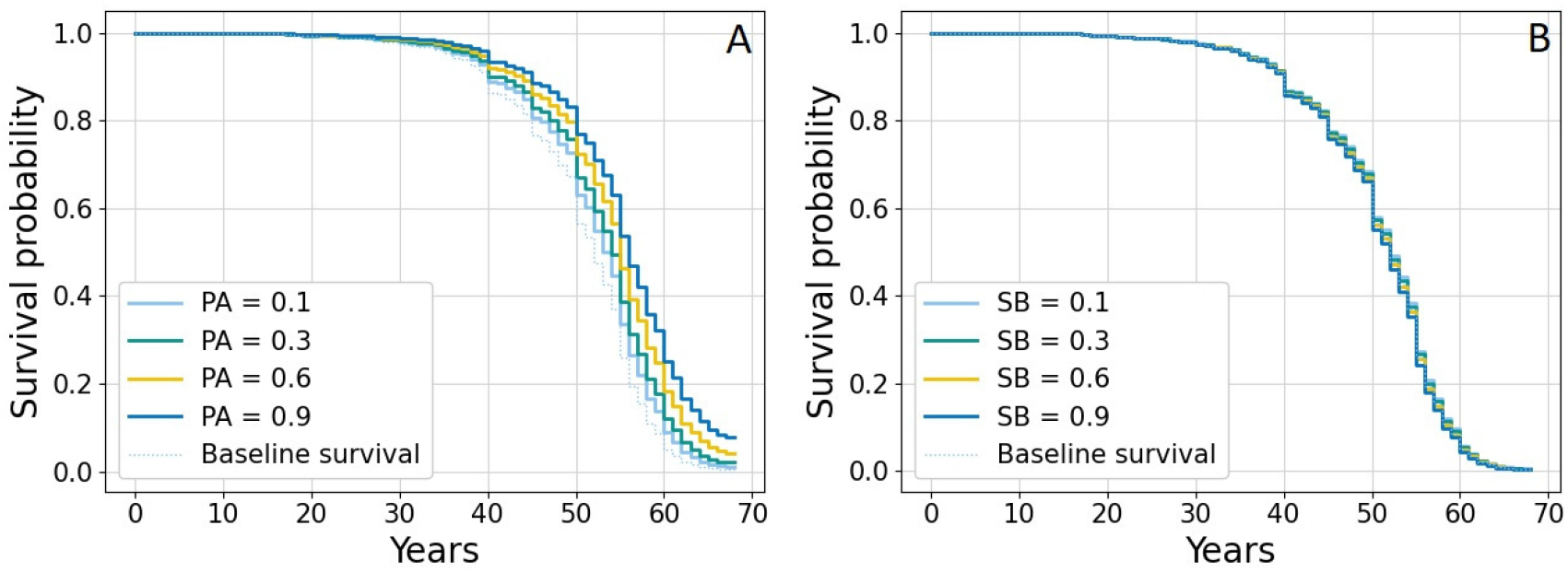
Partial effects of PA (A) and SB (B) on the survival function.

**TABLE VII:** Cox’s proportional hazards model prediction on censored subjects. Data presented as mean (standard deviation).

| Cluster | $n_{train}$ | $n_{young}$ | Predicted survival for *young* set |
| --- | --- | --- | --- |
| 1 | 365 | 143 | 19 (11) |
| 2 | 306 | 80 | 23 (12) |
| 3 | 314 | 64 | 20 (9) |
| 4 | 230 | 61 | 21 (13) |

## VI. Discussion

In this paper, we presented a new knowledge-based system for personalised activity recommendations. The system aims to increase PA and reduce SB in people with T2D by providing recommendations with a high chance of compliance based on the patient’s T2D cluster, previous activity, genetic risk and leptin levels. We trained and simulated the system using data from the UK Biobank and demonstrated that the system can help increase levels of activity. In fact, simulating the system with different probabilities of adherence showed improved physical activity compared to real-life activity measured after 1.5 years from baseline.

The novelty of the presented work lies in the adoption of a holistic approach to behaviour change interventions: in fact, we used a wide array of data—including biomarkers, genotype, leptin levels and past activity—to achieve highly personalised recommendations. A novel concept in our work is the use of genetic information to optimise the personalised recommendations. Genetic risk communication has been previously used for lifestyle behaviour change interventions, but has been found to have a very small effect on reducing unhealthy behaviours such as smoking, physical activity and nutrition [66]. However, to our knowledge, no intervention has been developed that uses genetic markers associated with the target behaviour. The inclusion of such markers may contribute to improving lifestyle behaviours because, in our system, genetic information is related to the capability of the subject. On the other hand, previous interventions have mapped genetic information to motivation and specifically to the knowledge of the risks associated to a poor lifestyle. Future research should confirm whether genetic associations can be used to directly inform lifestyle recommendations and achieve a significant change in lifestyle behaviours.

### A. Limitations of the current work

This work presents several limitations. Firstly, the behavioural analysis is limited as it only employed findings from a literature review, while conducting new qualitative research with the target population could have provided further insights into barriers and facilitators of the target behaviour, specifically in regard to mHealth interventions. Nonetheless, it is possible to improve the behavioural analysis by conducting focus groups or semi-structured interviews with the target population and incorporating the results in future versions of the proposed system.

Secondly, the system is evaluated through computer simulations rather than through a randomised controlled trial, which is the gold standard to assess the efficacy of an intervention [67]. A direct consequence is the impossibility of implementing features, such as goal customisation, which require the user’s input. Additionally, at this stage, it is not possible to assess the system’s user-friendliness and how it would impact the acceptance and efficacy of the intervention. It is also not possible to assess whether communicating the risk of developing CVD is the best way to increase motivation, and other rewards may be more effective. Additionally, several issues can undermine the efficacy of the system in a real-life setting, which could not be accounted for in simulations. Firstly, the engagement with the system is currently modelled with time-changing probability distributions, which may be different from the real-world adherence to the system’s recommendations. Secondly, modelling leptin levels can introduce errors in the optimisation of recommendations and thus in the efficacy simulations.

A further limitation of the current system version is the use of overall activity, which combines any activity not classified as sedentary time or sleep. Classifying activities based on type and intensity may be more beneficial for the user as it can provide a more precise indication of which kind of activity one should replace sedentary time with.

### B. Considerations for real-world implementation

The promising results presented in this paper warrant the implementation of the system into a real-life intervention, whose efficacy can be tested with a randomised controlled trial. To implement the intervention, the system would be integrated with a point-of-care test for leptin, which we presented in previous works [68], [15]. Additionally, the recommendations can be augmented with information on activity mode and duration, which are especially important when dealing with cardiometabolic conditions that require special considerations due to the associated risks. Additional gamification features, such as goal setting and scoreboards, can be integrated into the system to render the system customisable by the user and increase self-efficacy. As data availability increases and new features are added to the system, data can be represented using knowledge graphs to capture complex relationships and better integrate data from different sources. Specifically, a personal knowledge graph [69] would be used to define structural relationships between the patient and their data, which can then be used to improve recommendations, infer missing data, and predict disease risk.

However, before implementing the system into a real-world intervention, several considerations have to be made. While personalisation promotes acceptance of mHealth services [70], it also raises privacy and data security concerns. Tangari et al. [71] found that very few mHealth apps respect user privacy–88% of apps found in the major app stores can access and share personal data. As the present system relies on sensitive health data, its implementation must ensure privacy and user data security by using encrypted HTTP traffic and encrypting the data stored in the user’s device. Therefore, while mHealth applications often present a lack of data security, ensuring a high level of data protection is achievable in order to optimise the risk-benefit trade-off of mHealth interventions.

Once the initial implementation of the mHealth system is completed according to the best data protection practices, a first round of user testing should be conducted to assess the system’s usability, perceived benefits, and prospective acceptability. Subsequently, the system should be updated to reflect the feedback collected during the user trial. Finally, a randomised controlled trial would assess the efficacy of the proposed system, which is a first attempt at providing comprehensive and highly personalised activity recommendations by linking sensing data with genetics and daily tracking. If proved effective and acceptable in real-life settings, implementation of such theories could have a major impact in the prevention and management of chronic diseases that are triggered by behavioural practices.

## Funding

FRC is supported by the UK EPSRC Doctoral Training Partnership (Award Reference 2124497).

## Data Availability

The datasets analysed for this study can be obtained from the UK Biobank with appropriate application (ukbiobank.ac.uk).

## Notes

### Competing Interest Statement

CT is the co-founder of DnaNudge.

