## Supplementary information for "A knowledge-based system for personalised lifestyle recommendations: Design and simulation of potential effectiveness on the UK Biobank data"

### Supplementary Material

#### 1 Supplementary Data

##### 1.1 Normal ranges for biomarkers

- Age at onset > 45 years
- BMI < 30 kg/m2
- Waist circumference < 102 cm for males, 88 for females
- Glucose < 5.6 mmol/L
- HbA1c < 42 mmol/mol
- Systolic BP < 140 mmHg
- Diastolic BP < 90 mmHg
- Pulse rate 85-145 bpm
- Apolipoprotein A > 1.2 g/L
- Apolipoprotein B < 1 g/L
- Lipoprotein A < 75 nmol/L
- Triglycerides < 1.7 mmol/L
- Cholesterol < 5.2 mmol/L
- LDL direct < 2.6 mmol/L
- HLD-cholesterol > 1 mmol/L
- C-reactive protein < 2 mg/L

##### 1.2 List of diseases within the CVD classification

Heart/cardiac problem, peripheral vascular disease, venous thromboembolic disease, angina, heart attack/myocardial infarction, heart failure/pulmonary oedema, heart arrhythmia, heart valve problem/heart murmur, cardiomyopathy, pericardial problem, stroke, transient ischaemic attack, subdural haemorrhage/haematoma, subarachnoid haemorrhage, leg claudication/ intermittent claudication, arterial embolism, deep venous thrombosis (DVT), pulmonary embolism +/- DVT, cerebral aneurysm, myocarditis, atrial fibrillation, rheumatic fever, atrial flutter, wolff parkinson white/WPW syndrome, irregular heartbeat, sick sinus syndrome, supraventricular tachycardia, brain haemorrhage, aortic aneurysm, other venous/lymphatic disease, varicose veins, lymphoedema, ischaemic stroke, mitral valve disease, aortic valve disease, hypertrophic cardiomyopathy, pericarditis, pericardial effusion, aortic aneurysm rupture, aortic dissection, varicose ulcer, mitral valve prolapses, mitral stenosis, mitral regurgitation/incompetence, aortic stenosis.

##### 1.3 Ovid search strategy for review on barriers and facilitators to physical activity and sedentary behaviour

1) (physical train* or physical activ* or physical endur*).ti.
2) (sedentary behavio* or sedentary time* or sedentary lifestyle* or ((break* or interrupt*) adj3 sedentar* adj3 time*)).ti.
3) (barrier* or limitat* or imped* or restrict* or difficult*).ti.

4) (facilitat* or enable* or ease or assist*).ti.
5) ((type 2 diabetes or insulin-resistant diabetes or adult-onset diabetes or non-insulin dependent diabetes) not gestational not child*).ti.
6) 1 or 2
7) 3 or 4
8) 5 and 6 and 7
9) Remove duplicates from 8
10) ((behavio* change or intervention* or plan or program* or polic* or approach* or scheme or framework or guideline or concept) not trial).ti.
11) 9 and 10
12) Remove duplicates from 11

#### 2 Supplementary Tables

**Supplementary Table 1.** Summary of selected studies assessing barriers and facilitators to physical activity and sedentary behaviour in people with type 2 diabetes

| | | | | | |
|---|---|---|---|---|---|
| **Title** | Identifying barriers and facilitators to diet and physical activity behaviour change in type 2 diabetes using a design probe methodology | Understanding physical activity facilitators and barriers during and following a supervised exercise programme in Type 2 diabetes: a qualitative study. | Barriers to Physical Activity in People With Type 2 Diabetes Enrolled in a Worksite Diabetes Disease Management Program. | Barriers and enabling factors to use of a physical activity behavioural intervention for adults with Type 2 diabetes delivered in primary care: qualitative findings from the Movement as Medicine open pilot study: P519. | Frequency of Diet and Physical Activity Goal Attainment and Barriers Encountered Among Adults With Type 2 Diabetes During a Telephone Coaching Intervention. |
| **Authors** | K A Cradock et al. | D Casey et al. | D Erickson et al. | L Avery et al. | C M Swoboda et al. |
| **Doi** | https://doi.org/10.3390/jpm11020072 | 10.1111/j.1464-5491.2009.02873.x | https://doi.org/10.1177/0145721713492565 | https://doi.org/10.1111/DME.12668_1 | 10.2337/cd17-0023 |
| **Country in which the study conducted** | Ireland | Canada | United States | UK | United States |
| **Aim of study** | To understand the perceived barriers and facilitators to diet and PA behaviour change in persons with T2D. | To assess exercise facilitators and barriers in type 2 diabetes during and following a supervised exercise programme. | The purpose of this study was to explore the level of physical activity, barriers to physical activity, and strategies used to meet physical activity goals in people with T2DM. | To explore the barriers and enabling factors to patient acceptability of a behavioural intervention targeting physical activity behaviour of adults with type 2 diabetes. | 1) What are the types of dietary and PA goals selected, and how frequently is each goal type selected and attained? 2) How frequently and for how long do participants maintain a goal after attaining it the first time? and 3) What are common barriers to attaining |

| | | | | | |
|---|---|---|---|---|---|
| | | | | | self-set dietary and PA goals? |
| **Study design** | | Qualitative research | Cross sectional study | Qualitative research | Randomised controlled trial |
| **Population description** | | Participants that had engaged in a supervised exercise programme. | | 14 non-insulin dependent adults with type 2 diabetes. | |
| **Inclusion criteria** | Have T2D | Participants that had completed final assessment and had at least done one exercise class. | Participation in the worksite diabetes disease management program for 3 months or longer; diagnosis of T2DM; age at least 18 years; ability to speak, comprehend, and write in the English language; absence of a serious illness that would impair participants from engaging in moderate-intensity physical activity; and ability to give informed consent. | | Overweight or obese, 40-75 yrs old, with T2d for at least 1 year, have at least one additional risk factor for CVD, |
| **Exclusion criteria** | | | Residence in a facility where the individual was not responsible for self-management of diabetes. | | T1D or gestational diabetes, pregnancy, BMI > 50, medical issues requiring dietary treatment, inability to perform PA, clinical depression |

| | | | | | |
|---|---|---|---|---|---|
| **Total number of participants** | 21 | 16 | 75 | 14 | 37 |
| **Barriers and facilitators** | Physical health, mental health, social support, motivation, energy, weather, work schedule | Sustaining motivation, tangible health benefits, consciousness of facts not being enough to motivate, staying off diabetes medications is motivating, absence of supervision, supervision and support, supervision and support, fear of not having a support system, weather conditions, health concerns associated to diabetes, weather conditions, other commitments and time constraints | Having to change diabetes management as a consequence of increased PA, physical capability, time constraints, cost of equipment, availability of suitable environment | Detailed planning, knowledge about PA as a management option for T2D, initial high cognitive demand, active participation process, feedback on impact of PA on glycaemic control, feedback on impact of PA on glycaemic control, increased self-efficacy about ability to achieve PA goals | Time management, physical limitations, short term illness, feeling overwhelmed or stressed, weather, limited environmental resources, major life events |
